# The immune landscape of the inflamed joint defined by spectral flow cytometry

**DOI:** 10.1101/2023.11.30.569010

**Authors:** Meryl H. Attrill, Diana Shinko, Vicky Alexiou, Melissa Kartawinata, CHARMS study, JIAP study, Lucy R. Wedderburn, Anne M. Pesenacker

**Affiliations:** Institute of Immunity and Transplantation, Division of Infection and Immunity, University College London, London, UK; UCL Great Ormond Street Institute of Child Health, Infection, Immunity, and Inflammation Research and Teaching Department, University College London, London, UK; Centre for Adolescent Rheumatology Versus Arthritis at UCL UCLH and GOSH, London, UK; Centre for Rheumatology, Division of Medicine, University College London, London, UK; NIHR Biomedical Research Centre at GOSH, London, UK

**Keywords:** spectral flow cytometry, Juvenile Idiopathic Arthritis, synovial fluid, cellular adaptations, immune cell composition, inflammation

## Abstract

Cellular phenotype and function are altered in different microenvironments. For targeted therapies it is important to understand site-specific cellular adaptations. Juvenile Idiopathic Arthritis (JIA) is characterised by joint inflammation, with frequent inadequate treatment responses. To comprehensively assess the inflammatory immune landscape, we designed a 37-parameter spectral flow cytometry panel delineating mononuclear cells from JIA synovial fluid (SF), compared to JIA and healthy control blood. Synovial monocytes and NK cells lack the Fc-receptor CD16, suggesting antibody-mediated targeting may be ineffective. B cells and DCs, both in small frequencies in SF, undergo maturation with high 4-1BB, CD71, CD39 expression, supporting T cell activation. SF effector and regulatory T cells were highly active with newly described co-receptor combinations that may alter function, and suggestion of metabolic reprogramming via CD71, TNFR2 and PD-1. Most SF effector phenotypes, as well as an identified CD4-Foxp3+ T cell population, were restricted to the inflamed joint, yet specific SF-predominant Treg (CD4+Foxp3+) subpopulations were increased in blood of active but not inactive JIA, suggesting possible recirculation and loss of immunoregulation at distal sites. This first comprehensive dataset of the site-specific inflammatory landscape at protein level will inform functional studies and the development of targeted therapeutics to restore immunoregulatory balance and achieve remission in JIA.

**Graphical Abstract:** 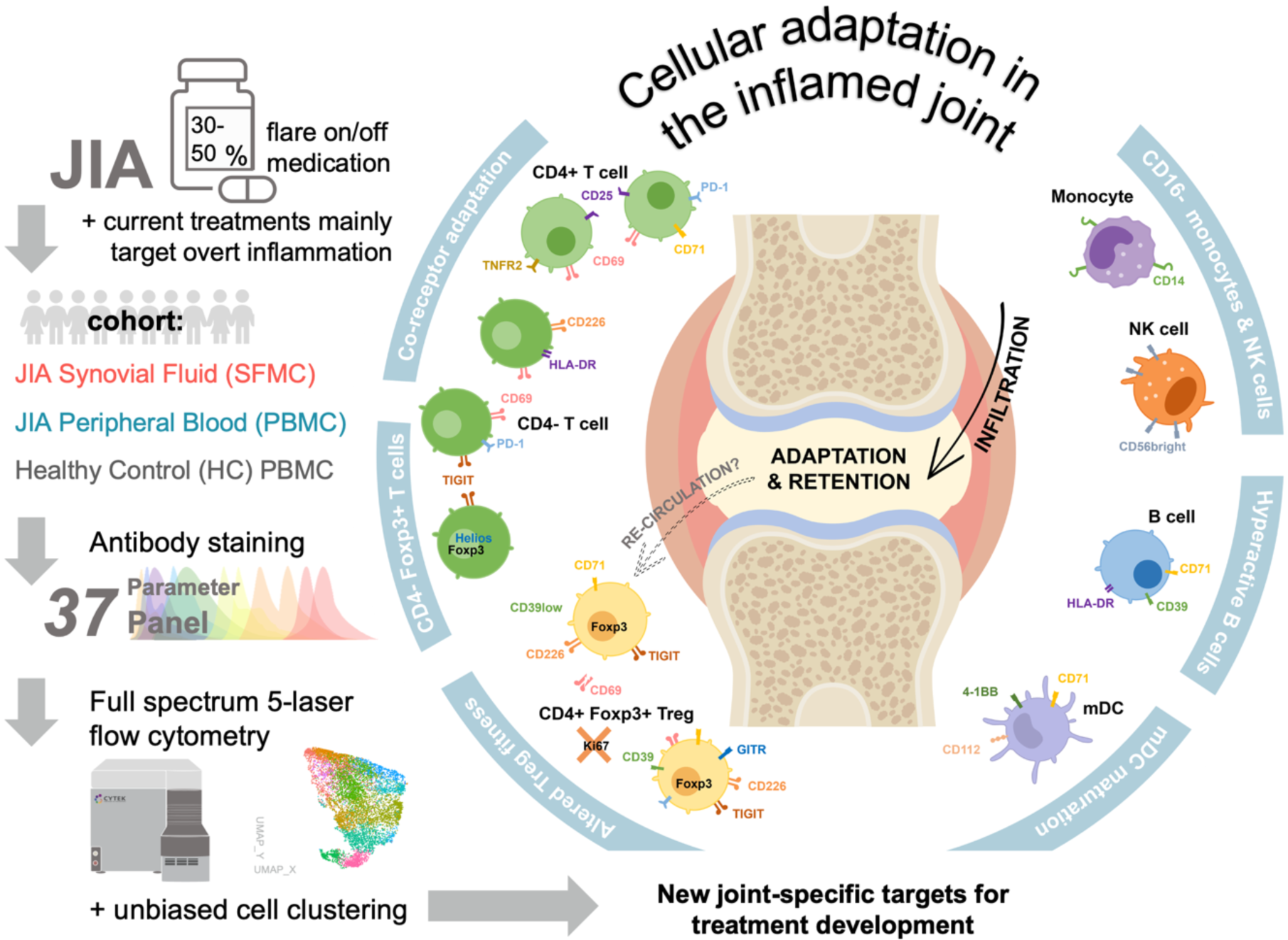

## Introduction

Juvenile Idiopathic Arthritis (JIA) is the most common autoimmune rheumatic disease of childhood-onset, affecting over two million people worldwide (1, 2). JIA is defined as the sustained presence of arthritis of unknown cause with the onset before the age of 16, with current ILAR classification categorising seven subtypes differing in clinical features, additional organ involvement and genetic susceptibility (1, 3). Here we focus on oligoarticular (persistent or extended) and polyarticular Rheumatoid Factor (RF) negative JIA, the most common subtypes accounting for 40-60% of all JIA cases and characterised by joint inflammation without skin, ligament or systemic involvement (1, 4). Repeated inflammatory flares without warning or known trigger lead to pain, loss of mobility, reduced quality of life and ultimately joint destruction and disability (1).

Current treatment approaches for JIA are focused on targeting the symptoms of overt inflammation, with a stepwise approach taken based mainly on the clinician’s experience and preference (1, 5). Treatments range from general immunosuppression by corticosteroids (oral and/or local joint injection), methotrexate and targeted biologics (e.g. TNF-α blockade) (1, 5). Whilst these treatments have undoubtedly improved the lives of many children and young people with JIA, they can have severe systemic side effects including nausea, increased risks of infections, and potential growth disturbances (1). Additionally, 30-50% of patients do not achieve adequate responses, experience flares on treatment or treatment inefficiency over time (1, 5). Moreover, disease activity can worsen with each flare, increasing the chance more joints affected (6). Along with an overactive immune system, regulatory T cell (Treg) dysfunction is a hallmark of JIA, yet therapeutics targeting this regulatory arm are currently lacking (1, 7). Understanding the cellular adaptations and unique immune landscape of the inflamed microenvironment, and identifying any pathogenic cells in recirculation, are key hurdles to therapeutically alter the local environment, restore immunoregulatory balance and achieve better treatment outcomes.

The precise etiology of JIA is still largely unknown. Although autoantibodies such as anti-nuclear antibodies are common in oligoarticular and polyarticular JIA, their involvement in disease pathogenesis is limited (1, 4). Instead, immune infiltration and increased pro-inflammatory cytokines drive joint inflammation with Tregs unable to control this process (1, 4, 7, 8). In conjunction with corticosteroid injection, synovial fluid (SF) can be aspirated and analysed, showing an enrichment in T cells (70-90% of immune cells), impaired neutrophils, and highly activated memory lymphocytes (8–12). Interestingly, CD4+Foxp3+ Tregs are also enriched in the inflamed joint. SF Tregs retain hypomethylation of the TSDR (Treg specific demethylated region), showing commitment to the Treg fate (13), and decreased numbers might associate with disease severity (12). However, several genetic risk alleles in JIA are in loci associated with Treg function (7), SF Tregs can exhibit a pro-inflammatory effector phenotype, loss of IL-2 sensitivity and their *in vitro* suppressive capacity has been questioned in some studies (5, 12–20). Thus, the effects of the pro-inflammatory microenvironment on SF immune cells, including Tregs, need to be further assessed to understand their role in ongoing disease. Moreover, previous studies of the SF immune composition have focused on either the major immune populations or assessing one specific subset, thus a comprehensive analysis of the immune network with detailed phenotype is missing and is required to understand the pathogenesis of JIA. We therefore aim, in this study, to fully investigate the cellular changes in composition and phenotype in the inflamed joint of JIA to assess the overall immunoregulatory balance and identify key drivers of inflammation.

The advancement of spectral flow cytometry has enabled the development of high-dimensional immunophenotyping at single-cell protein level even in small samples (21, 22). Here, we developed a 37-parameter panel utilising a 5-laser full spectrum cytometer and unbiased clustering approaches to highlight microenvironmental adaptations across cell types in SF from the inflamed joint of individuals with JIA and peripheral blood (PB) from JIA and healthy controls. Markers for activation, functionality, and adaptations through co-receptor expression could identify cell subsets unique to active joint inflammation by comparing populations in SF and PB samples from oligoarticular/RF-polyarticular JIA. We identified immune cell subsets of each lineage that were highly activated, matured, with markers of adaptations, and restricted to the inflamed joint. Our deep phenotyping of SF and PB of active and clinically inactive JIA (by active joint count, AJC) revealed retention of effector and antigen presenting cells (APCs) within the inflamed joint but potentially dysfunctional Treg subsets recirculating through blood of patients with active but not inactive disease. These identified phenotypes driving continued inflammation in JIA may help to understand disease pathogenesis and be suitable as new therapeutic targets specific to the inflamed site without systemic immune suppression. Conversely, targeting identified pathways which may enable the ‘resetting’ of the immunoregulatory balance may lead to more individuals in sustained remission.

## Results

With the introduction of spectral flow cytometry, it is now possible to extensively map the broad immune cell landscape as well as detailed phenotypic definition of cell adaptations with small clinical isolates. Here, we utilised 5-laser spectral flow cytometry for high-dimensional single cell level analysis of JIA synovial fluid (SFMCs, n=18), compared to PBMCs isolated from individuals with JIA (n=52) and healthy control (HC, n=18, age range 22-43, mean 28). JIA samples were a cohort of oligoarticular (69.1%) or RF-polyarticular (30.9%) JIA, with an age range of 1-18 (mean 8.3) and predominantly female (78.6%) and Caucasian (71.4%) (Table 1).

**Table 1.** Sample demographics and clinical characteristics of JIA cohorts.

|  | HC PBMC | JIA PBMC | JIA SFMC |
| --- | --- | --- | --- |
| **Number of participants, n** | 18 | 52 | 18 |
| **Gender, % Female\*** | 50.0 | 75.0 | 88.9 |
| **Age at sample in years, mean (range) \*** | 28.6 (22-43) | 7.4 (1-18) | 11.0 (4-15) |
| **Ethnicity, % Caucasian \*** | 78.6 | 63.5 | 72.2 |
| **Disease duration:** |  |  |  |
| **Time since diagnosis, months (range) $\diamond$** | N/A | 37.0 (2-125) | 92.5 (19-158) |
| **JIA subtype:** |  |  |  |
| **% RF- Polyarticular** | N/A | 34.0 | 22.2 |
| **% Oligoarticular** | N/A | 66.0 | 77.8 |
| **Medication at time of sample:** |  |  |  |
| **Methotrexate, n $\Delta$** | N/A | Yes= 27<br>No= 20 | Yes = 4<br>No = 7 |
| **Steroids, n $\nabla$** | N/A | Yes= 24<br>No= 22 | Yes= 4<br>No= 7 |
| **Biologics, n $+$** | N/A | Yes= 4<br>No= 42 | Yes= 1<br>No= 7 |
| **Clinical information at time of sample:** |  |  |  |
| **ANA, n $\#$** | N/A | positive = 29<br>negative = 12 | positive = 14<br>negative = 2 |
| **AJC, mean (range) $\blacklozenge$** | N/A | 1.28 (0-5) | 1.82 (1-4) |
| **cJADAS, mean (range) $\wedge$** | N/A | 5.63 (0-17.3) | 6.50 (2.4-13.8) |

PBMC= Peripheral Blood Mononuclear Cells; SF= Synovial Fluid; HC= Healthy Control; RF-: Rheumatoid factor negative; ANA: Antinuclear Antibodies (positive classified as titre ≥ 1:160); AJC: Active Joint Count; cJADAS: clinical Juvenile Arthritis Disease Activity Score(102)
\* 22.2% of HC PBMC samples missing data; $\diamond$ 7.7% of JIA PBMC samples missing data; $\Delta$ 9.6% of JIA PBMC and 38.9% of JIA SFMC samples missing data; $\nabla$ 11.5% of JIA PBMC and 38.9% of JIA SFMC samples missing data; $+$ 11.5% JIA PBMC and 55.6% of JIA SFMC samples missing data; $\#$ 21.1% of JIA PBMC and 11.1% of JIA SFMC samples missing data; $\blacklozenge$ 11.5% of JIA PBMC and 38.9% of JIA SFMC samples missing data; $\wedge$ 23.1% of JIA PBMC and 55.6% of JIA SFMC samples missing data

A 37-parameter panel (Table S1) was developed to simultaneously enumerate the frequencies of myeloid, B, NK, T cell and Treg subpopulations alongside detailed phenotyping of activation status, proliferative capacity, expression of known immunoregulatory markers and co-receptor expression profile, with the aim of identifying local microenvironmental cellular adaptations at the site of inflammation.

### The inflamed joint cellular composition consists of increased CD4- T cell and CD4+Foxp3+ Treg populations, with decreased B cell frequencies compared to blood

Unbiased FlowSOM clustering (23) on concatenated live HC PBMCs, JIA PBMCs and JIA SFMCs identified 18 cellular metaclusters of myeloid, pDC, B, NK, and T cell subsets defined by 32 different markers (Figure 1A-C). The clusters identified were confirmed by assessing known lineage markers and traditional gating strategies (Figure 1C). T cells were defined by CD3+, with CD4+ conventional T cells (clusters 1,5), Foxp3+CD4+ Tregs (clusters 6,7), and CD4- T cells (clusters 2,8). B cells were categorised by CD19 expression (clusters 10,14,18), NK and NKT cells by CD56+ and CD3+CD56+ respectively (cluster 3,4), pDCs by high HLA-DR and CD123 expression (cluster 13), and additional myeloid cell subsets by CD11c expression (clusters 9,12,15-17). A minor cell population in cluster 11 could not be attributed to any specific linage and was therefore excluded from further analyses.

**Figure 1.**
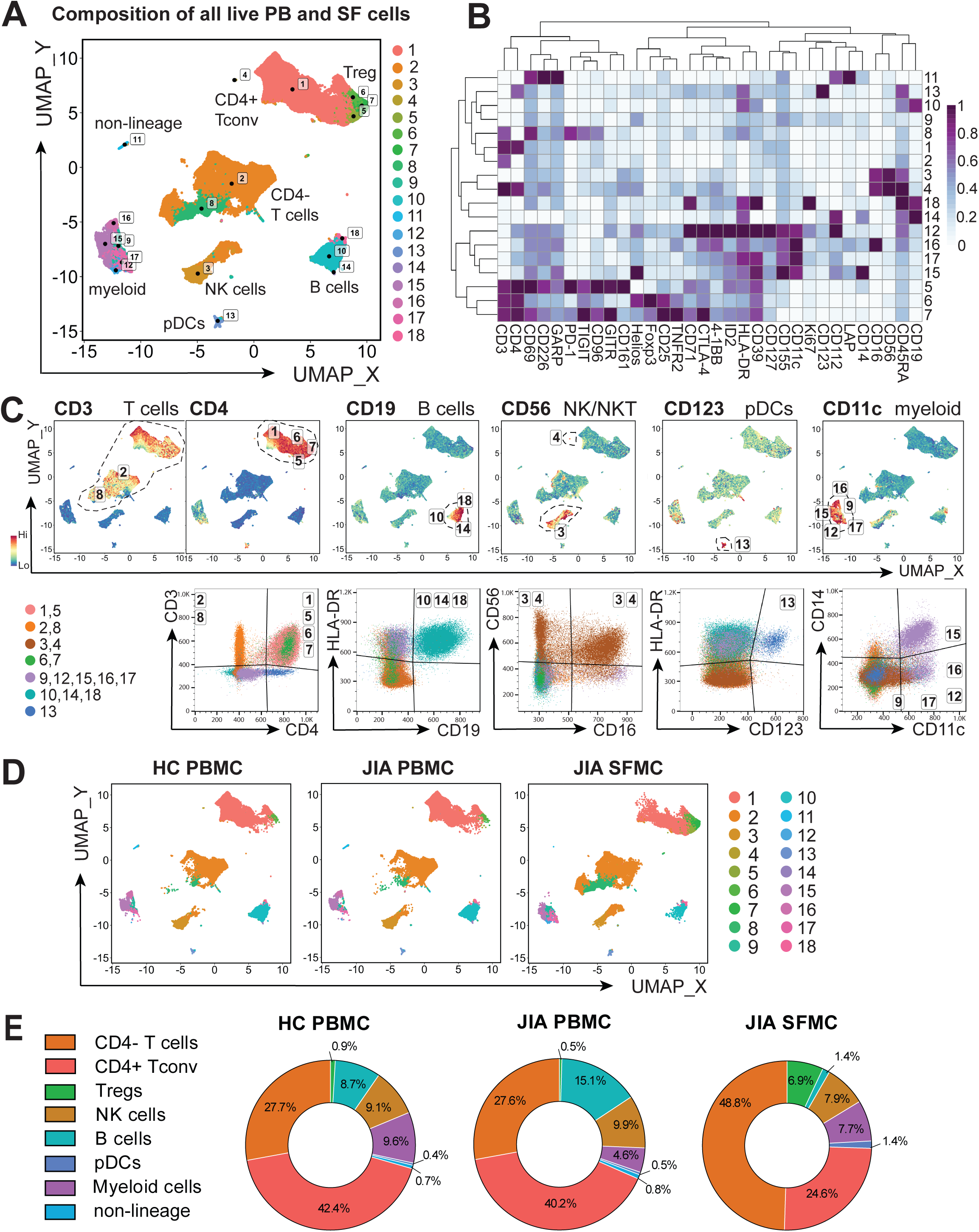
Cell composition is altered in the inflamed joint of JIA. Synovial fluid mononuclear cells (SFMC, n=18) and PBMC of JIA patients (n=52) and healthy controls (HC, n=18) were stained and data were acquired on a full-spectrum cytometer. Data were analysed using an unbiased clustering algorithm FlowSOM, after gating on all single, live (FVD-) cells. (**A**) UMAP identifying the 18 clusters of the major cellular compositions in all samples. (**B**) Heatmap of the 32 markers used for clustering, Z-score across cluster rows determined by median fluorescence intensity (MFI). (**C**) Expression UMAP plots of lineage and phenotypic markers CD3, CD4, CD19, CD56, CD123, and CD11c and validation of the cluster identity using flow plots. (**D**) UMAP of the 18 clusters in HC PBMC, JIA PBMC, and JIA SFMC. (**E**) Frequencies of the major immune cell subpopulations utilising cell type assignment of Figure 1C and a normalised UMAP of all samples per sample group in HC PBMC, JIA PBMC, and JC PBMC.

These parent cell populations were present in PB and SF, but at variable frequencies (Figure 1D-E). Gating on the eight cellular cluster identities confirmed that SF was highly enriched in T cells (80.3% CD3+ of live SFMCs) with an expected increase of CD4- T cell populations compared to PB (48.8% CD4- of live SFMC vs 27.7% HC PBMC and 27.6% JIA PBMC, Figure 1E) (8). CD4+Foxp3+ Treg frequencies were additionally 7.7- and 13.8-fold higher in SF compared to HC and JIA PB respectively (6.9% of live SFMCs vs 0.9% HC PBMC and 0.5% JIA PBMC, Figure 1E).

While T cells were enriched, B cell frequency was decreased 10-fold in the inflamed joint (1.4% of live SFMC vs 8.7% HC PBMC and 15.1% JIA PBMC, Figure 1E), as shown previously (8). The frequency of CD56+ NK cell populations varied little between groups (9.1%, 9.9%, 7.9% for HC PBMC, JIA PBMC, JIA SFMC respectively, Figure 1E). Myeloid cell frequency consisted of one small cluster of pDCs (0.4%, 0.5%, 1.4% of HC PBMC, JIA PBMC and JIA SFMC respectively, Figure 1E) and five additional myeloid subset clusters (combined total of 9.6%, 4.6%, 7.7% of all HC PBMC, JIA PBMC and JIA SFMC respectively, Figure 1E).

### Distinct myeloid subpopulations primed for survival are enriched at the inflamed site

While the total myeloid frequency was consistently below 10% of all live mononuclear cells across SF and PB (Figure 1E,2A), five different CD11c+ subsets (clusters 9,12,15, 16 and 17, Figures 1-2) demonstrated distinct phenotypic diversity. Four of these myeloid populations differed in frequency between PB and SF (clusters 12,15,16 and 17). In contrast, cluster 9 myeloid cells with lower CD11c expression were not significantly different in frequency between blood and SF (Figure 2B-C).

**Figure 2.**
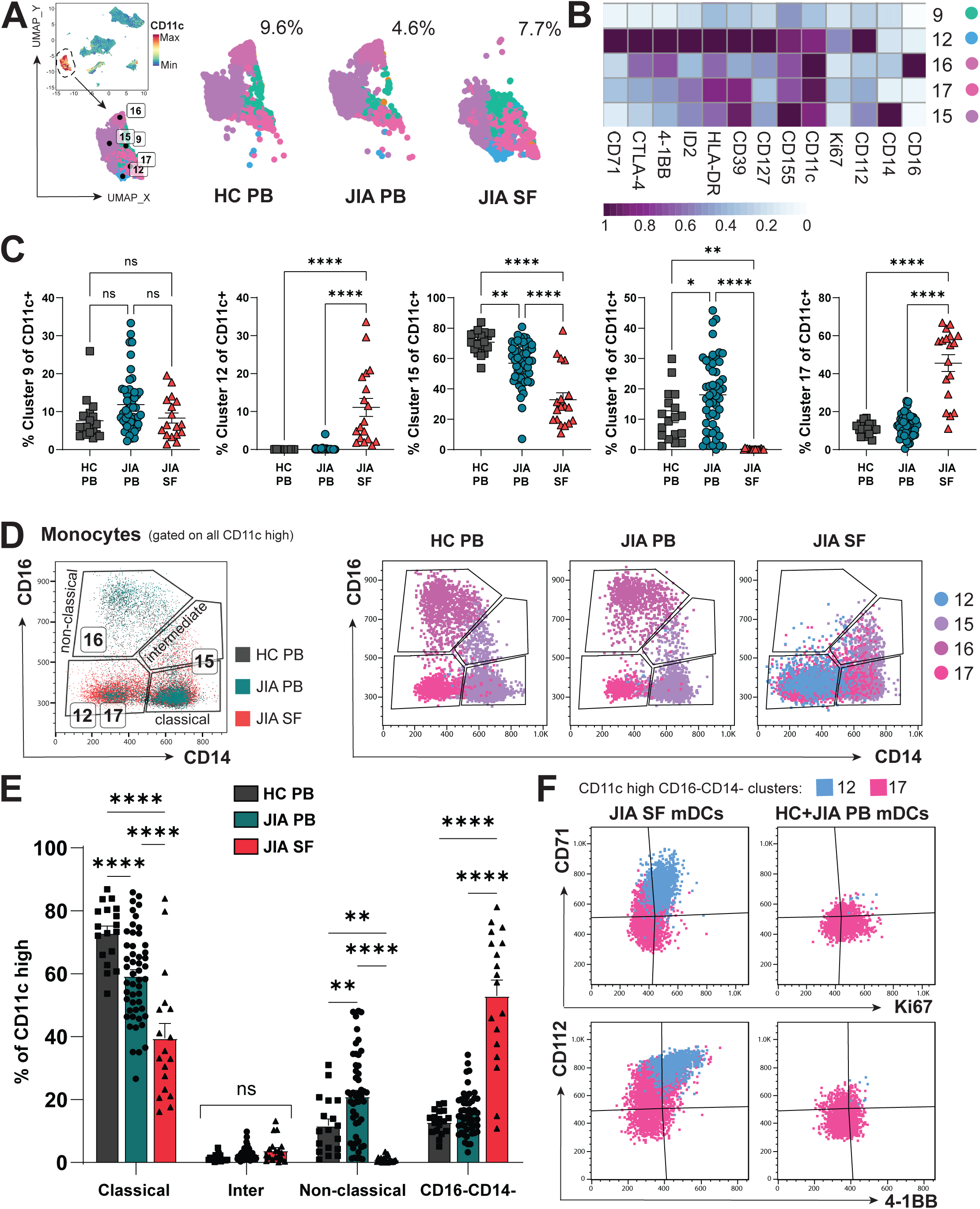
The inflamed joint may promote adaptation and survival of distinct myeloid cells. Data were gated on single, live cells and analysed using unbiased clustering algorithm FlowSOM. (**A**) Expression UMAP of CD11c identifying five myeloid clusters (9, 12, 15, 16, 17) and UMAP of myeloid clusters in HC PB, JIA PB, and JIA SF, with % CD11c+ clusters of all live cells shown per sample group. (**B**) Heatmap of myeloid clusters with differentially expressed markers, Z-score across cluster rows. (**C**) Frequencies of myeloid clusters 9, 12, 15, 16, 17 between HC PB, JIA PB, and JIA SF as percentage of all CD11c+ myeloid clusters. (**D**) *Left:* Representative flow plot (pre-gated on CD11c high) showing classical, intermediate, and non-classical monocyte gating by CD14 and CD16 expression and the location of the myeloid clusters 12, 15, 16, and 17 within the manual gating. *Right:* Flow plots of the myeloid clusters (12, 15, 16, 17) and monocyte gating across HC PB, JIA PB, and JIA SF. (**E**) Frequencies of the gated classical, intermediate, non-classical monocytes, and CD16-CD14- cells (as % of CD11c high) in HC PB, JIA PB, and JIA SF. (**F**) Flow plots of phenotypic difference displaying CD71, Ki67, CD112, and 4-1BB in the monocytic dendritic cells (mDCs; CD11c+CD14-CD16-) clusters 12 and 17 between JIA SF and JIA+HC PB. Throughout: JIA Synovial fluid cells (SF; n=18), JIA peripheral blood cells (PB; n=52), healthy control PB (HC; n=18). Data are represented as mean±SEM with parametric two-way ANOVA with Tukey’s multiple comparison testing, ** p<0.01, **** p<0.0001, ns= non-significant.

Cluster 16 containing a population of CD16lowCD14high cells represent pro-inflammatory non-classical monocytes (24), which were at a low frequency or undetected in SF but a large monocyte population observed in blood (0.14±0.04% of SF CD11c+ cells vs HC PB 10.8±1.9% and JIA PB 18.0±1.6%, mean±SEM, Figure 2C-D). Cluster 15, which encompassed both intermediate (CD16intermediateCD14+) and classical (CD16-CD14+) monocytes (Figure 2D), was reduced in SF (32.8±4.6% of SF CD11c+ cells) compared to HC and JIA PB (70.7±1.7%, 56.9±1.9% respectively, p<0.0001). This difference in monocyte phenotype frequencies identified by unbiased clustering were additionally confirmed through manual gating of CD14 and CD16 (SF 0.95±0.23% CD14lowCD16high of CD11c+ myeloid cells vs HC PB 11.7±2.1%, p=0.0088 and JIA PB 21.0±1.9%, p<0.0001, Figure 2D-E). Whilst there was no statistical difference in the frequency of intermediate monocytes in the inflamed joint compared to PB, classical monocytes were significantly reduced in number (39.5±4.7% CD16-CD14+ of SF CD11c+ cells vs HC PB 72.9±2.2% or JIA PB 59.3±2.0%, p<0.0001, Figure 2E), thus driving reduction of cluster 15. The absence of non-classical and significant reduction of classical monocytes in the inflamed joint, compared to blood, suggests a potential abrogation of non-classical monocyte-mediated resolution of inflammation (25).

The majority of SF CD11c+ myeloid cells, unlike PB, were encompassed by cluster 17 (45.6%±4.4) and cluster 12 (11.1%±2.4), expressing neither CD14 nor CD16 with higher HLA-DR levels, likely representing myeloid dendritic cells (mDCs) (53.0±5.0% of SF CD11c+ myeloid cells vs 12.9±0.9% HC PB, 15.7±1.0% JIA PB, p<0.0001, Figure 2E). Interestingly, cluster 12 was unique to the inflamed joint environment (Figure 2C) and had a proliferating, activated phenotype with high Ki67, CD71, 4-1BB (CD137) and HLA-DR expression (Figure 2B,F). CD71 expression was also increased in mDCs of Cluster 17 present in the inflamed joint but not in cluster 17 cells from blood (Figure 2F). Increased CD71 expression might indicate metabolic adaptation to the inflammatory environment by this transferrin receptor, which enables essential iron supply, promoting cell metabolism and growth (26). Similarly, co-receptor ligand CD112 (nectin-2) and co-stimulatory receptor 4-1BB were highly expressed on SF mDC clusters 12 and 17, but not PB mDCs from the same clusters (Figure 2B,F). 4-1BB has been described to increase the survival, longevity and subsequently immunogenicity of dendritic cells (27), whilst CD112 expression has been associated with increased DC maturation, acting as a ligand for TIGIT or CD226 (DNAM-1) (28).

Taken together, this suggests synovial mDCs are adapted in the inflamed, metabolically restricted microenvironment with a profile indicative of enhanced maturation, activation, proliferation, and survival. Moreover, SF in JIA appears devoid of non-classical monocytes and has significantly reduced proportions of classical monocytes compared to PB, thus SF myeloid populations are skewed away from resolution-linked phenotypes.

### Synovial B cell proportions are reduced but show a hyperactive phenotype

As we confirmed, B cell frequency in the inflamed joint is dramatically decreased compared to PB (Figure 3A)(8, 10). Despite this, it has been suggested that B cell hyperactivity might contribute to JIA pathogenesis (29). Indeed, B cell cluster 18, which was found almost exclusively in the joint (47.3±5.5% of SF B cells vs 3.1±1.4% of HC PB and 2.7±0.3% of JIA PB B cells, Figure 3D), were characterised by high CD71 expression (Figure 3B-D), a transferrin receptor upregulated on activated lymphocytes. Interestingly, while the few cluster 18 B cells found in PB expressed high levels of proliferation marker Ki-67, SF cluster 18 B cells were largely Ki-67 negative (Figure 3D), suggesting that although highly active, these SF B cells are non-proliferative. Co-receptor ligand CD112 and HLA-DR expression characterised B cell cluster 14 across all sites (Figure 3E), even though it was a low frequency cluster in PB (0.54±0.04% of HC PB, 0.59±0.04% of JIA PB B cells) and near absent in SF (0.14±0.04% of all SF B cells). Cluster 10 B cells, which were dominant in both HC and JIA PB (96.3±1.4%/96.7±0.3% of HC/JIA PB B cells vs 52.6±5.5% of SF B cells, p<0.0001, Figure 3F), corresponded to a more classic resting, circulating B cell phenotype of CD71- CD112- Ki67-. However, SF cluster 10 B cells showed some expression of CD71 and CD69 (Figure 3B,F), potentially indicating recent activation in the inflamed joint without yet having transitioned to a different cluster phenotype.

**Figure 3.**
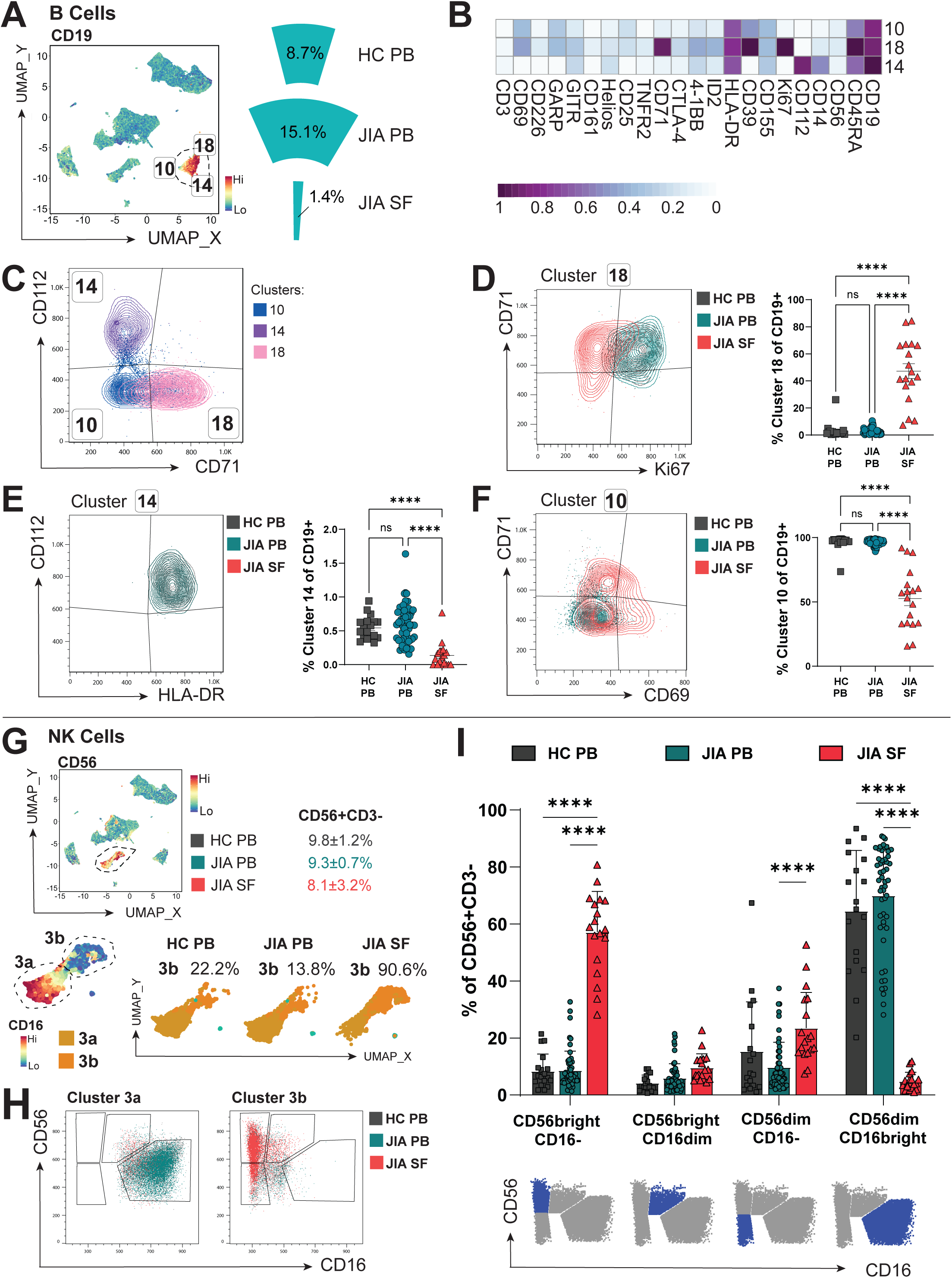
JIA SF has distinct B and NK cell populations from PB. Data were gated on single, live cells and analysed using an unbiased clustering algorithm, FlowSOM. (**A**) Expression UMAP of CD19 identifying three B cells clusters (10, 14, 18) and frequencies of all B cell clusters across HC PB, JIA PB, and JIA SF (as % of total live cells) (**B**) Heatmap of B cell clusters and differentially expressed markers, Z-score across cluster rows. (**C**) Overlay flow plot displaying the phenotypic differences separating the B cell clusters 10, 14, and 18 using CD112 and CD71. (**D-F**) *Left:* Flow plots displaying CD71, Ki67, CD112, HLA-DR, and CD69 illustrating the phenotype and *right:* frequencies (of % of CD19+ B cells) of B cell clusters 18 (D), 14 (E), and 10 (F) across HC PB (grey), JIA PB (blue), and JIA SF (pink). (**G**) *Top:* CD56 expression UMAP identifying NK cell cluster 3 and the frequencies of CD56+CD3- cells across HC PB, JIA PB, and JIA SF. *Bottom:* Subclassification of NK cluster 3 (3a and 3b) based on CD16 expression UMAP and the frequencies of cluster 3b in HC PB, JIA PB, and JIA SF. (**H**) Gating strategy for NK cells subtypes and phenotypic difference between cluster 3a and 3b based on CD56 and CD16 expression, with the relative expression across HC PB, JIA PB, and JIA SF shown. (**I**) Frequencies of NK cell subtypes (as % of CD56+CD3-) in HC PB, JIA PB, and JIA SF and the gating location of the gated subpopulation (blue) in the CD56 vs CD16 flow plots. Throughout: JIA Synovial fluid cells (SF; n=18), JIA peripheral blood cells (PB; n=52), healthy control PB (HC; n=18). Data are represented as mean±SEM with two-way ANOVA with Tukey’s multiple comparison testing, **** p<0.0001.

### Synovial NK cell subpopulations are distinct from PB

Through unbiased clustering and enumerating CD56+CD3- NK cells, we found total NK cell numbers were not significantly altered in the inflamed joint compared to PB (%CD56+CD3- of all live cells 8.1±3.2 in SF vs 9.8±1.2% HC PB, 9.3±0.7% JIA PB, Figure 3G). However, SF and PB CD56+CD3- cells were grouped in separate subclusters (denoted as Cluster 3a and 3b, Figure 3G). Traditional gating showed that while CD56dimCD16bright NK cells were the predominant NK population in PB (encompassed by cluster 3a, HC PB 64.6±5.0%, JIA PB 70.0±2.5%), this population was near absent in SF (SF 4.8±0.7%, Figure 3H-I). In contrast, SF predominated in cluster 3b CD56brightCD16- NK cells (57.1±3.4% of CD56+CD3- NK cells vs HC PB 8.4±1.4%, JIA PB 8.7±0.9%, Figure 3H-I). Thus, SF NK cells, just as SF monocytes, appear to have lost CD16 protein expression indicating that antibody-mediated targeting may be ineffective. Whether this is due to shedding, CD16 being completely occupied by autoantibodies in the inflamed joint, or if to alter NK/monocyte functionality remains to be investigated. CD56brightCD16- NK cells, dominant in SF, have increased cytokine production and reduced cytotoxic capabilities than CD56dim counterparts and therefore likely contribute to the inflammatory milieu (30–32).

### Highly activated T cell subsets adapt co-receptor expression in the inflammatory microenvironment

The greatest differences in cellular composition between PB and SF were within CD3+ T lymphocytes, which made up the majority of mononuclear cells (69.6±1.9% HC PB, 67.7±1.2% JIA PB, 77.5±1.6% JIA SF, Figure 4A) but with differing distribution of CD4+ and CD4- T cells. After gating on CD3+CD19- (Figure 4A) for CD3 T cell sub-clustering by FlowSOM, T cells segregated into two distinct areas via UMAP: CD3i and CD3ii (Figure 4B). CD3ii T cells were largely CD4+ while CD3i defined CD4- cells, representing predominantly CD8+ T cells, likely with a small proportion of gamma-delta T cells (8). Indeed, we confirmed an inversion of the CD4+:CD4- T cell ratio within the SF compared to blood (SF 0.90±0.16, HC PB 1.75±0.15, JIA PB 1.63±0.07, Figure 4C), as previously suggested (8). SF CD3+ T cell clusters were consistently differentiated from PB by high CD69 expression (Figure 4D), suggesting selective recruitment, recent activation within the inflammatory environment (33) or indicating tissue retention in the joint (34, 35).

**Figure 4.**
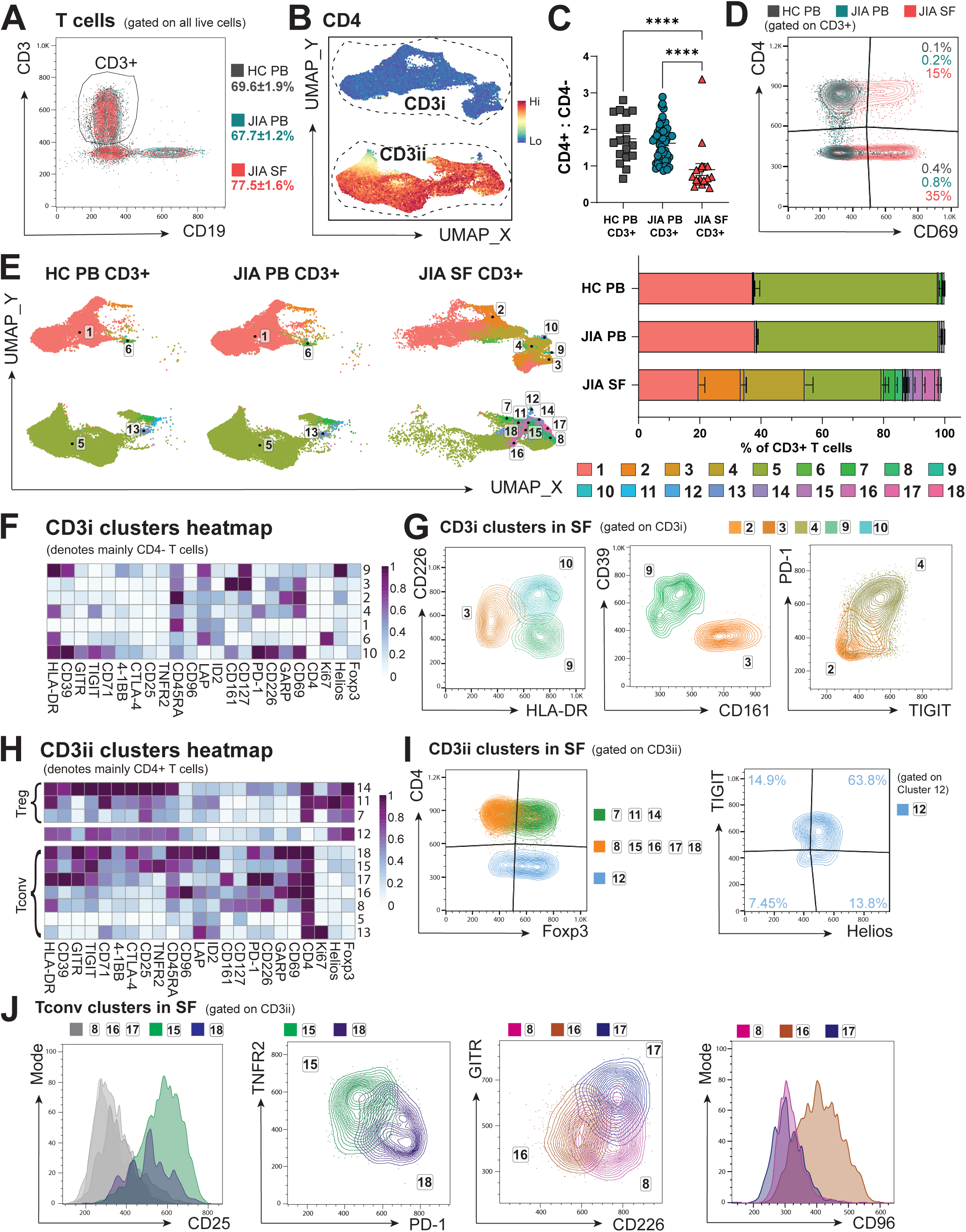
T cell subsets are highly activated and adapt co-receptor expression in the inflammatory SF microenvironment. Data were gated on single, live, CD3+CD19- cells and sub-clustered using unbiased clustering algorithm, FlowSOM. (**A**) Gating and frequencies of CD3+ T cells (as % of live cells) in HC PB, JIA PB, and JIA SF. (**B**) Subclassification of CD3 clusters into CD3i and CD3ii using CD4 expression UMAP. (**C**) Ratio of CD4+ to CD4- T cells (as % of CD3+) in HC PB, JIA PB, and JIA SF. (**D**) Flow plot showing CD69 expression across HC PB, JIA PB, and JIA SF CD3+ T cells. (**E**) *Left:* UMAP plots and *right:* frequencies (as % of CD3+ T cells) of the 18 T cell clusters in HC PB, JIA PB, and JIA SF. (**F**) Heatmap of the CD3i clusters (denoting mainly CD4- T cells) and their expression of relevant T cell markers, Z-score across cluster rows. (**G**) Overlay flow plots displaying the phenotype of CD3i clusters (2, 3, 4, 9, and 10) in SF using CD226, HLA-DR, CD39, CD161, PD-1, and TIGIT. (**H**) Heatmap of the CD3ii clusters (denoting mainly CD4+ T cells) split into Foxp3+ regulatory T cells (Treg) clusters and FoxP3- conventional T cells (Tconv) clusters and their expression of relevant T cell markers, Z-score across cluster rows. (**I-J**) Overlay flow plots displaying the phenotype of CD3ii clusters in SF (7, 8, 11, 12, 14–18) with (I) showing Foxp3 across CD3ii clusters and cluster 12 phenotype using TIGIT and Helios, and (J) phenotype of the CD3ii CD4+ Tconv clusters in SF with flow plots displaying CD25, TNFR2, PD-1, GITR, CD226, and CD96. Throughout: JIA Synovial fluid cells (SF; n=18), JIA peripheral blood cells (PB; n=52), healthy control PB (HC; n=18). Data are represented as mean±SEM with one-way ANOVA with Tukey’s multiple comparison testing, **** p<0.0001.

Unbiased clustering on CD3+ SF and PB T cells identified 18 distinct T cell clusters, utilising 23 cell markers (Figure 4E,F,H). Most PB T cells were encompassed by clusters 1 and 5 (97.6%±0.24 of HC PB T cells, 97.6%±0.25 of JIA PB T cells vs 41.1%±3.18 of JIA SF T cells, Figure 4E), representing a PB- dominant resting circulating phenotype for CD4- and CD4+ T cells respectively (Figure 4F,H). However, T cell clusters 1 and 5 in SF had increased levels of CD69 compared to the same clusters in PB and SF cluster 5 cells also expressed higher levels of CD71, HLA-DR and PD-1. This may suggest that the SF T cells of cluster 1 and 5 cells had recently infiltrated the inflamed joint, began to adapt but had not differentiated sufficiently to cluster separately. Other PB CD3+ T cells were divided into proliferating Ki67+ CD4- T cells (cluster 6, 0.39±0.08% of HC PB, 0.60±0.10% of JIA PB, 0.93±0.20% of SF CD3+, Figure 4E,F) and Ki67+ CD4+ conventional T cells (Tconv, cluster 13, 0.56±0.10% of HC PB CD3+, 0.64±0.08% of JIA PB CD3+, 0.58±0.11% of SF CD3+, Figure 4E,H). Interestingly, as the only significant Ki67+ clusters not differing between SF and PB, SF T cells were overall no more proliferative than PB, despite increase in activation markers such as CD69.

Out of the 18 CD3+ clusters, 14 were significantly SF-predominant, displaying considerable T cell heterogeneity in the inflammatory environment (Figure 4E). Within the CD3i cluster group, the SF- dominant CD4- clusters 2, 3, 4, 9, and 10 were discernible by co-stimulatory co-receptor CD226, co-inhibitory co-receptors PD-1 and TIGIT, and activation marker HLA-DR (Figure 4E-G). Cluster 10 likely represents a small fraction of highly activated cells with high CD226 and HLA-DR expression (0.5%±0.24 of SF T cells, 0.0% in HC/JIA PB, Figure 4G). CD161highCD226intermediateHLA-DRlow cluster 3 cells may represent CD4- mucosal associated invariant T (MAIT) cells (1.26%±0.49 of SF, 0.0% of HC/JIA PB T cells, Figure 4F-G), which have been reported to be enriched in different tissues (36–38). Cluster 9 denotes a small fraction of T cells with high expression of HLA-DR and CD39 but low CD226 expression (0.04%±0.21 of SF, 0.0% of HC/JIA PB T cells, Figure 4E,G). The expression of CD45RA but low expression of PD-1 (CD279), TIGIT and other activation markers besides CD69 suggests cluster 2 CD4- cells (13.8%±1.6 of SF vs 0.28%±0.03 of HC PB and 0.49%±0.06 of JIA PB T cells, Figure 4E,G) could represent TEMRA (terminally differentiated effector memory cells) that re-express CD45RA (39).

The greatest proportion of SF-prominent CD4- T cells were in cluster 4 (19.6±2.6% of SF CD3+ vs 0.27%±0.05 of HC PB and 0.27%±0.07 of JIA PB T cells, Figure 4E). These had a phenotype of HLA- DRhighPD-1+TIGIT+ (Figure 4G) suggesting a largely exhausted CD4- T cell phenotype in SF (40, 41). Alternatively, increased PD-1 expression has been shown to drive T cell metabolic adaptation (42) and thus this phenotype could allow these cells to survive within the inflamed joint.

The CD3ii group of T cell clusters mainly encompassed CD4+ T cells (Figure 4B, H-J), with the exception of cluster 12 which was specific to the inflamed joint (0.23±0.05% of SF CD3+ T cells vs 0.0% of HC/JIA PB CD3+ T cells, Figure 4E). This CD4- cluster largely expressed Foxp3, Helios and TIGIT (Figure 4I) as well as CD25, CD39, CTLA-4 (CD152), CD71, HLA-DR and low CD127 (Figure 4H), thus mirroring CD8 Treg-like cells which have been described in inflamed joints of rheumatoid arthritis (RA), psoriatic arthritis (PsA) and spondylarthritis (43, 44). Accordingly, cluster 12 was grouped in close proximity to CD4+ Treg clusters 7,11 and 14 (Figure 4E), highlighting their phenotypic similarities (44). Conversely, Foxp3 expression by cytotoxic CD8 T cells might aid a metabolic shift and survival, similar to that seen in the tumour microenvironment (45).

CD4+ Tconv clusters 8, 15, 16, 17 and 18 were only detected in synovial fluid (0.0% of HC/JIA PB CD3+, Figure 4E). These Tconv clusters could be distinguished by expression of CD25, TNFR2 (CD120b), PD-1, CD226 and GITR (CD357) (Figure 4J). CD4+ Tconv clusters 15 (3.03±0.58% of SF T cells) and 18 (0.50±0.18% of SF T cells) were marked by high CD25 expression (Figure 4J), denoting activated populations (46, 47). These CD25high Tconv clusters were separated by TNFR2 and PD-1 expression, with cluster 18 potentially exhibiting a more exhausted phenotype (48) of PD-1high TNFR2low, and cluster 15 TNFR2highPD-1low (Figure 4J). TNFR2 expression could mediate a strong response to TNF-α in the SF and resistance to Treg suppression, as demonstrated in mice (49, 50). Indeed, TNF-α-dependent mechanism has been suggested for JIA SF effector T cell resistance (14, 15). PD-1 and TNFR2 engagement on CD4+ T cells have also been implicated in enabling metabolic flexibility upon TCR activation and therefore could promote survival within the inflammatory environment (42, 51).

Of the CD25low CD4+ Tconv clusters, cluster 17 was characterised by high GITR (a TNFR-family receptor) and co-stimulatory co-receptor CD226 expression (1.25±0.59% of SF T cells, Figure 4E, J), suggesting a small but previously activated co-stimulation responsive population (52). Cluster 8 (2.40±0.70% of SF T cells) expressed high CD226 but low GITR, whereas cluster 16 (3.99±0.83% of SF T cells) was low/intermediate for both CD226 and GITR but expressed the highest level of co-receptor CD96 (Figure 4H, J). CD96 competes for CD155 with TIGIT and CD226 but its function on T cells remains unclear (53).

Thus, SF CD4+ and CD4- conventional/effector T cells were generally activated with phenotypes for potential metabolic adaptations that may enhance survival in the inflammatory environment through CD71, PD-1, TNFR2 and/or Foxp3 upregulation, with differences in co-receptors and specific activation markers.

### Treg fitness is altered in the inflamed joint

Since total T cell clustering showed heterogeneity within CD4+Foxp3+ clusters, we gated on CD3+CD4+Foxp3+ cells (Figure 5A) and applied PhenoGraph clustering algorithm to further investigate Treg phenotypes across PB and SF (Figure 5B-G). We defined 15 phenotypically distinct Treg clusters across 20 markers (Figure 5B-D). As expected, Tregs were significantly increased in frequency in SF (19.4±1.6% of SF CD4+CD3+ cells vs 4.4±0.4% of HC PB and 4.3±0.3% of JIA PB CD3+CD4+ cells, Figure 5A), but in addition, five Treg clusters (4, 7, 9, 10, 11) were limited to SF, while Treg clusters 1, 5, 8, 12, 13, 14 predominated in PB (Figure 5B-C). As shown in Figure 5E, the SF-predominant Treg clusters had high expression of activation markers (CD69, CD71, GITR, 4-1BB and HLA-DR), markers associated with ‘effective’ Treg suppressive functionality (CD39, TNFR2, CTLA- 4, TIGIT), and co-receptors able to alter TCR signalling outcomes (TIGIT, CD226, CD96, PD-1, 4-1BB). Overall, Treg ‘fitness’ can therefore be assessed by the varied expression of these markers and co-receptors, altering signalling or metabolism, and thus functional capacity, of Tregs at the inflamed site.

**Figure 5.**
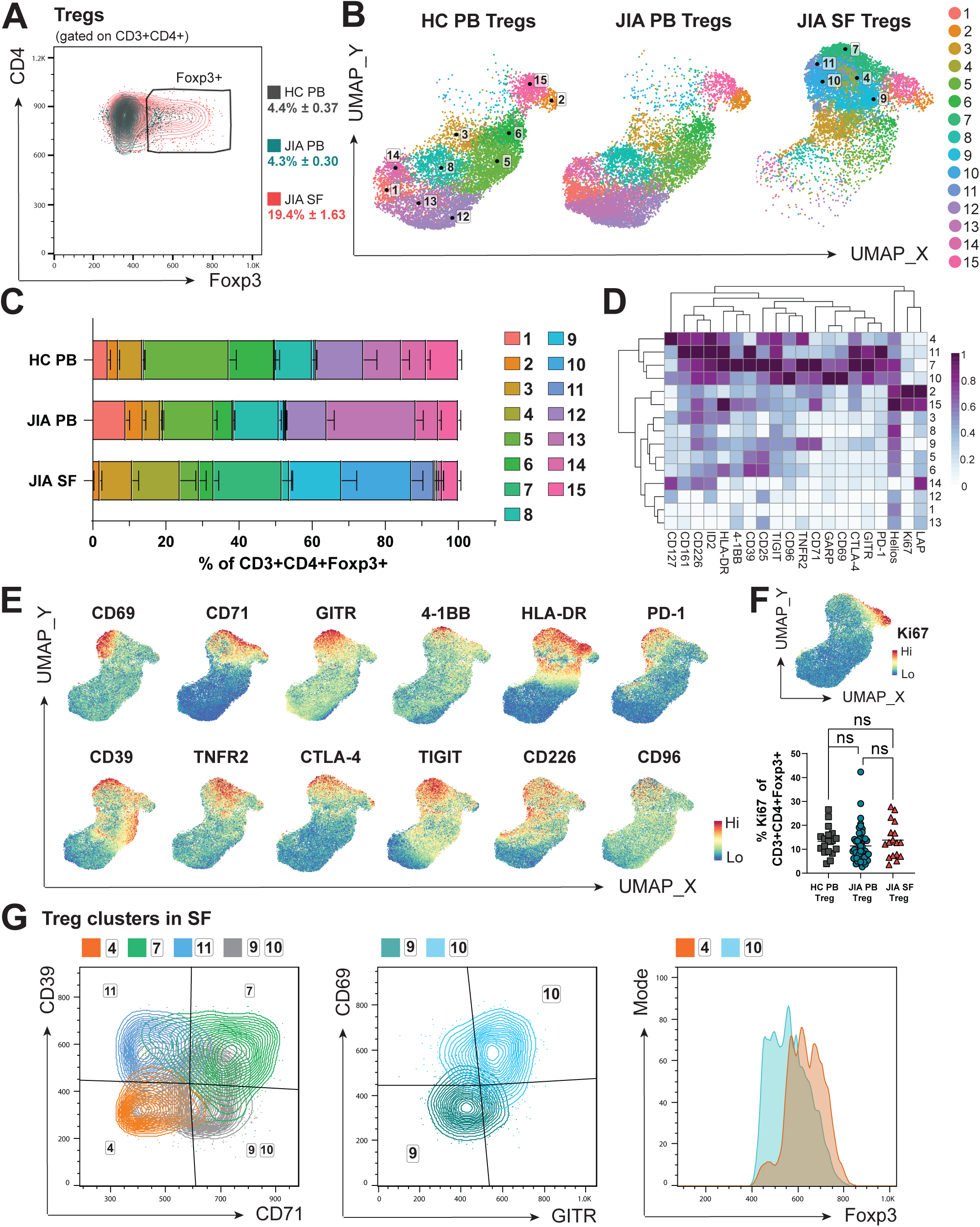
Regulatory T cell fitness is altered in the inflamed JIA joint. Data were gated on single, live, CD3+CD4+Foxp3+ regulatory T cells (Tregs) and sub-clustered using unbiased clustering algorithm, PhenoGraph. (**A**) Gating strategy and frequency of Foxp3+ cells (as % of CD3+CD4+) in HC PB, JIA PB, and JIA SF. (**B**) UMAP of the 15 Treg clusters in HC PB, JIA PB, and JIA SF. (**C**) Frequencies of the 15 Treg clusters (as % of CD3+CD4+Foxp3+) in HC PB, JIA PB, and JIA SF. (**D**) Heatmap of the Treg clusters and their expression of the 20 relevant Treg markers used for clustering, Z-score across cluster rows. (**E**) UMAP of combined HC PB, JIA PB and SF PB Tregs with heatmap overlay of expression levels (by median fluorescent intensity) of specific markers found in SF-predominant clusters. (**F**) UMAP of all groups combined showing Ki67 expression as heatmap overlay with summary plot for Ki67 frequency (as % of CD3+CD4+Foxp3+) in HC PB, JIA PB, and JIA SF. (**G**) Overlay flow plots displaying the phenotype of Treg clusters 1, 7, 9-11 in SF using CD39, CD71, CD69, GITR, and Foxp3. Throughout: JIA Synovial fluid cells (SF; n=18), JIA peripheral blood cells (PB; n=52), healthy control PB (HC; n=18). Data are represented as mean±SEM with one-way ANOVA with Tukey’s multiple comparison testing, ns = not significant.

Clusters 2 and 15 were clearly separated from other clusters by high Ki67 expression, denoting cycling Tregs, which were present across all sites (Cluster 2: 2.82±0.28% of HC PB Tregs, 4.73±0.74% of JIA PB Tregs, 1.80±0.51% of JIA SF Tregs; Cluster 15: 8.74±0.91% of HC PB, 5.40±0.61% of JIA PB, 4.38±0.69% of JIA SF Tregs, Figure 5B-C). Indeed, when analysing %Ki67+ of all Tregs, no differences were seen between sample types (Figure 5F). Therefore, the majority of SF Tregs are not proliferating despite increased activation, suggesting that the increased frequency of Tregs at the inflamed site is more likely due to increased recruitment and/or retention and survival.

The nature of SF-predominant Treg clusters could be further delineated by expression levels of CD39, CD71, CD69, GITR and Foxp3 (Figure 5D,G). The low CD71 and CD39 but increased CD127 expression and high Foxp3 levels of cluster 4 cells (9.6±2.73% of SF Treg, Figure 5C-D,G) suggests a resting Treg population with potentially decreased suppressive abilities (54, 55). Cluster 11 (6.17±2.52% of SF Treg) and 7 (18.7±2.89% of SF Treg) were associated with high expression of functional Treg markers including CD39 and CTLA-4 but differentiated in CD71 levels (Figure 5D,G). CD71high cluster 7 therefore likely denotes more activated Tregs and increased transferrin receptor CD71 may enable cells to utilise iron metabolism in response to the inflammatory environment (46, 56). High CD71 expression, but low CD39 levels were hallmarks of SF Treg clusters 9 (14.3±4.07% of SF Treg) and 10 (19.1±3.09% of SF Treg), but with differential expression of CD69 and GITR (cluster 9 CD69lowGITRlow, cluster 10 CD69highGITRhigh, Figure 5G). With high CD69 expression, cluster 10 may be more likely to be retained within the SF or have been more recently activated (35, 46). Cluster 10 could also represent SF Tregs that have lost sensitivity for IL-2 despite CD25 expression, with lower Foxp3 and high PD-1 levels (Figure 5D,G) as proposed by Bending et al (13). High PD-1 levels of clusters 7, 10 and 11, which together make up over 40% of SF Tregs (Figure 5C-D), could also demonstrate a metabolic adaptation of Tregs to allow for better survival in the inflamed environment (42) or hinder their functionality (57, 58).

Taken together these data demonstrate highly activated, non-proliferating Tregs in the inflamed joint. It is possible these distinct phenotypic profiles enable these Tregs to cope with the inflammatory environment through possible metabolic adaptation. However, despite expressing many functional Treg markers, SF Tregs are unable to control inflammation in JIA, potentially due to an imbalance in co-receptors altering their functionality.

### SF-predominant effector T cell and APC populations are mostly restricted to SF, but some SF Treg subsets can be detected in PB of active but not inactive JIA

To identify whether specific alterations in cell populations we detected in the SF could also be detected in blood of JIA individuals with active disease, and thus potentially represent re-circulating cells spreading inflammation across to distal joints, we next divided the cohort of JIA PB samples into clinically active (active joint count, AJC≥1, n=29) and inactive (AJC=0, n=17) to compare alteration driven by active disease in otherwise well-matched samples (Table S2). Interestingly, some statistically significant differences of clusters identified with SF samples were detected in these two JIA PB groups.

Myeloid cell clusters 15 (classic/intermediate monocytes) and 17 (activated mDCs) were reduced and increased respectively in active JIA PB compared to inactive PB (cluster 15, 60.9±2.4% of CD11c+ PB cells from inactive JIA vs 52.3±2.8% from active JIA PB samples, p=0.0278; cluster 17, 9.7±1.3% of CD11c+ PB cells from inactive JIA vs 13.5±1.1% from active JIA PB samples, p=0.0314). Active JIA PB therefore was closer to SF composition of these clusters than inactive PB was to SF. However, cells within cluster 17 show a different phenotype in PB compared to SF (see Figure 2F). SF cluster 17 cells may therefore originate from PB cluster 17 cells, but further activate after infiltration into the joint and are likely retained in SF.

On the other hand, while absent in SF, non-classic monocytes (cluster 16) were increased in active PB compared to inactive PB (14.5±2.9% inactive vs 21.7±2.1% active JIA, p=0.0496, Figure 6A compared to Figure 2C-E). Similarly, the CD16- NK phenotype detected in SF NK cells (see Figure 3I) were not mimicked in the periphery of active individuals. In contrast, CD56bright CD16- and CD56brightCD16dim NK cells were reduced, with more CD56dimCD16bright in active JIA PB compared to inactive JIA PB samples (CD56brightCD16- 13.2±2.28% inactive vs 5.77±0.46% active; CD56brightCD16dim 9.49±1.63% inactive vs 4.04±0.48% active; CD56dimCD16bright 60.2±5.40% inactive vs 77.4±2.02% active, Figure 6B). These data further indicate that the lack of CD16 is specific to the inflamed joint and might be due to shedding, receptor occupation or changes to functionality within the SF microenvironment.

**Figure 6.**
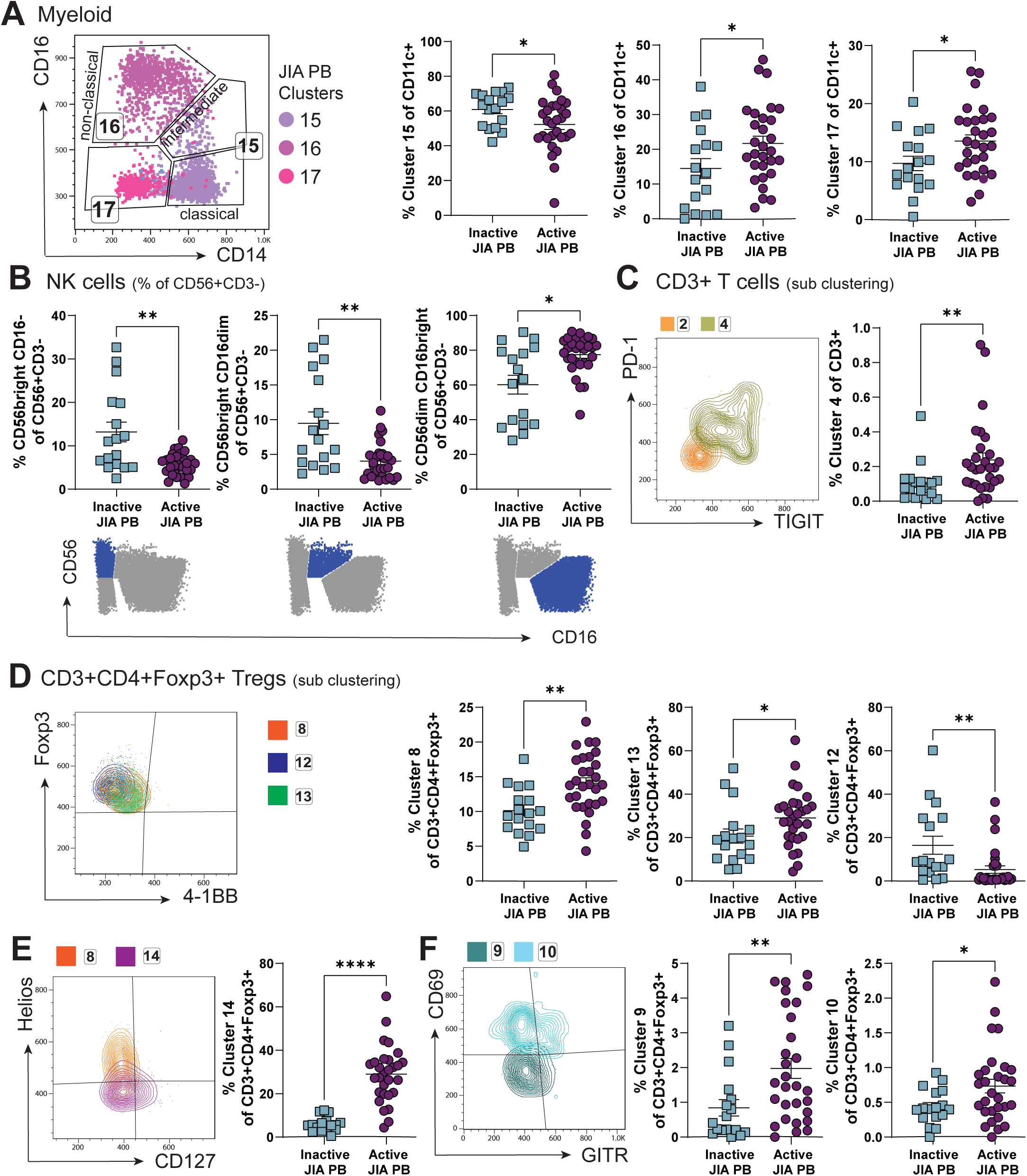
Unlikely recirculation of SF dominant clusters by assessing PB of active and inactive JIA. Clusters identified by studying JIA SF samples were assessed in clinically active (active joint count, AJC≥1) and clinically inactive (AJC=0) JIA PB samples, expecting to identify recirculating cells in active but not inactive JIA PB. (**A**) Gating (using CD14 and CD16 expression), phenotype, and frequency (as % of CD11c+) of myeloid clusters 15, 16, and 17 in JIA PB in inactive vs active JIA PB. (**B**) Frequencies (as % of CD56+CD3-) of NK cell subtypes by traditional gating strategy using CD56 and CD16 (symbolised below summary graphs) in inactive vs active JIA PB. (**C**) Overlay flow plot displaying the phenotype (using PD-1 and TIGIT) of clusters 2 and 4 from CD3+ T cell sub-clustering and frequency (as % of CD3+) of cluster 4 in inactive vs active JIA. (**D-F**) For sub-clustering of CD3+CD4+Foxp3+ Tregs, phenotype and frequencies (as % of CD3+CD4+Foxp3+) Treg specific clusters 8, 13, 12 (D), 14 (E), 9 and 10 (F) in inactive vs active JIA PB. Throughout: JIA peripheral blood cells (PB) of clinically inactive (AJC=0, n=17) and active (AJC≥1, n=29). Data are represented as mean±SEM with Mann-Whitney test, * p<0.05, ** p<0.01, **** p<0.0001.

CD4- T cells and CD4+ Tconv clusters that predominate in the SF were largely undetectable in PB (Figure 4E). Only T cell cluster 4 (CD4-PD-1highTIGIT+, Figure 4G, 6C) showed a statistical difference in PB when analysed against clinical activity, with increased frequency in PB samples from active compared to inactive JIA (0.11±0.03% of CD3+ in inactive JIA PB vs 0.24±0.04% of CD3+ in active JIA PB, p=0.0054, Figure 6C). Although the most prominent T cell cluster in SF, the frequency of cluster 4 in blood was very low, thus biological significance of the difference between active and inactive JIA PB samples is difficult to interpret. Overall, this suggests most CD3+ effector T cells that have infiltrated the inflamed joint adapt their phenotype significantly at the site and do not recirculate.

Interestingly, the most statistically significant differences between inactive and active PB samples were among CD3+CD4+Foxp3+ Treg clusters (Figure 6D-F). Although rare in SF, the resting PB-specific Treg clusters 8 and 13 were both at greater frequency in the blood of active JIA compared to inactive (cluster 8 10.1±0.79% of inactive vs 14.1±0.79% of active PB Tregs, p=0.0015; cluster 13 20.7±3.26% of inactive vs 29.0±2.43% of active PB Tregs, p=0.0191, Figure 6D). However, Cluster 12, another resting, memory Treg population which was also near absent in SF, was found to be decreased in the blood of individuals with active JIA compared to inactive (16.5±4.1% inactive vs 5.2±1.7% active JIA PB Tregs, p=0.0018). Whether some of these populations could be precursors of SF infiltrating Tregs remains to be seen.

Treg cluster 14 saw the largest difference in active compared to inactive JIA PB (29.0± 2.43 active vs 5.91± 0.80% inactive JIA PB Tregs, p<0.0001, Figure 6E). The high CD127 and low Helios expression on these cells could represent decreased suppressive function of Tregs in active disease (55) or represent activated conventional CD127high Tconv that have upregulated Foxp3 (59–62) and thus non-Treg contaminants in the Treg Foxp3 gate in PB, which did not see in the inflamed joint. This may suggest different activation states and function of Tconv in the periphery of JIA blood during an active inflammatory flare of the joint.

SF-dominant activated CD71highCD39low Treg clusters 9 and 10 were found at significantly higher frequencies in active compared to inactive JIA PB (cluster 9 0.84±0.23% of inactive vs 1.98±0.29% of active JIA PB Tregs, p=0.0055; cluster 10 0.44±0.06% of inactive vs 0.73±0.10% of active JIA PB Tregs, p=0.0401, Figure 6F). Interestingly, cluster 9 expressed low CD69 levels also in SF, thus could represent recirculating cells seen in the active PB samples. The expression of CD69 in cluster 10, with lower levels of Foxp3 expression could represent recently activated but latent Tregs with low CD39 (54) and potential loss of IL-2 sensitivity (13). Both these populations might circulate towards other joints and associated lymph nodes and there fail to stop activated mDCs promoting autoreactive T cell activation.

## Discussion

Cellular adaptation to changing microenvironments can enhance survival and alter functions within different niches. The phenotypic and metabolic adaptations of immune cells specific to the tumour microenvironment have been targeted therapeutically to improve anti-tumour immunity (63–66). The functional adaptation to inflammation within local tissues has been less well described with a recent drive to understand the inflammatory cellular adaptations and heterogeneity in rheumatic diseases (67). It is therefore important to consider the specific cellular adaptations that occur at the inflamed site in autoimmune conditions such as JIA, to enhance understanding of disease pathogenesis and improve cellular or molecular targeting.

Here, we utilised high-dimensional phenotyping at a single cell protein level by spectral flow cytometry for a broad and unbiased characterisation of the immune cell landscape (68) of the inflamed joint in JIA to improve understanding of the immune pathogenesis. Using a well characterised cohort of samples, we defined the cellular composition and heterogeneity between blood and synovial fluid of individuals with JIA, suggesting a distinct profile of immune cells infiltrate the joint and adapt to the microenvironment.

Here, we observe that synovial monocytes and NK cells lacked the expression of CD16, with almost a complete absence of CD14-CD16+ non-classical monocytes and CD56dimCD16bright NK cells, a similar profile to that seen in other inflammatory arthritis (69). Although Schmidt *et al.* observed a significant increase in intermediate CD14+CD16+ monocytes in SF of oligoarticular JIA (70). The absence of non-classical monocytes in the inflamed joint suggests a potential abrogation of non-classical monocyte-mediated resolution of inflammation (25). CD16 is an Fc receptor (FcγRIIIA) that is essential for antibody-mediated responses (71, 72), thus, SF NK and monocytes might be unable to perform antibody-mediated actions in the joint. This may suggest that treatments relying on CD16- antibody interactions may be ineffective. The lack of CD16 could be explained through shedding, which occurs post NK cell activation (73), or full occupation with Ig in the inflamed environment might block *ex vivo* CD16 staining.

In addition to a lack of CD16 expression, SF NK cells were predominately CD56bright, in line with previous studies (8, 69, 74). CD56bright NK cells have been associated with lower levels of perforin, granzymes, and cytolytic granules and increased pro-inflammatory cytokine production (30–32). Conversely, CD56bright NK cells may exhibit immunoregulatory functions (32, 75, 76), as shown in Multiple Sclerosis (75). In JIA, however, this suppressive function was found to be impaired (74). Thus, SF NK cells likely contribute to the pro-inflammatory microenvironment of SF by secretion of proinflammatory cytokines.

Moreover, SF-exclusive 4-1BB, CD71 and CD112 high mDC-like cells likely use these molecules for increased iron metabolism, survival and activation of T and NK cells (26–28). A restricted number of markers specific to B cells in inflammatory environments were included in this phenotyping panel, limiting comparisons to other studies in inflammatory disease (77). Nevertheless, we were able to confirm reduced B cell numbers in JIA SF (8), additionally displaying SF B cells to be highly activated, in agreement with the hyperactivity of B cells recently suggested as a possible pathogenesis factor of JIA (29, 78).

Highly activated monocyte, mDC and B cells in SF with altered co-receptor ligand expression, will affect the balance of signals delivered to T cells upon interaction. Indeed, SF T cell sub-clustering revealed high expression of multiple activation markers, whereas PB T cells were mainly resting. The transferrin receptor CD71 was also a common activation marker frequently found on SF T cells, especially on SF CD4+ and regulatory T cells. CD71 is essential for iron uptake via endocytosis of transferrin-bound iron, with its expression correlated to increased proliferation (79–81). However, despite increased CD71, the majority of SF T cells were not actively proliferating, demonstrated by the lack of Ki67 expression in SF-predominant clusters. Nevertheless, upregulation of CD71 in SF cells might represent a metabolic adaption to the inflamed joint, with increased iron uptake driving glucose metabolism through the mTOR signalling pathway (56). Alternatively, high levels of iron have been reported within synovial fluid of inflamed joints in RA (82), where iron overload may impair differentiation and function of Th1 and Th17, and death of Tregs through increased reactive oxygen species (79, 83). Th1, Th17 and Treg balance has been previously implicated in JIA pathogenesis (19, 84, 85). Similarly, elevated CD71 was found to alter T cell function, promoting a pro-inflammatory phenotype in lupus and idiopathic inflammatory myopathies (56, 86). Therefore, high expression of CD71 in JIA SF CD4+ and Treg subsets could be affecting the regulatory balance and, with further investigation, targeting iron metabolism could be a potential therapeutic approach in JIA.

Indeed, in this study multiple possible metabolic adaptions are suggested in SF T cells by change in marker expression. Besides being a marker associated with T cell exhaustion and reducing the level of T cell activation, PD-1 has also been linked to metabolic reprogramming from glycolysis to oxidative phosphorylation (40, 42, 87). We identified high PD-1 expression on two CD4-, four CD4+ T cell and three Treg subsets specific to the inflamed joint. While CD3+ cluster 4, with elevated TIGIT and fewer activation markers expressed, might represent an exhausted CD8 T cell population, the other PD-1+ SF T cell subsets express many activation and effector molecules. These may therefore represent metabolic and functional adaptation in response to local stimulation (48) with high proinflammatory capacity and cytokine production (88). PD-1-mediated shift to lipid metabolism can favour Treg function (42), thus indicating that the three PD-1+ SF Treg clusters making up over 40% of SF Treg clusters should be metabolically equipped to function. On the other hand, PD-1 can actively restrict Tregs as its blockade can lead to hyperprogressive cancers driven by activated Tregs (57, 58).

Conversely, recent data highlighted that TNFR2 (CD120b) ligation in Tregs leads to a shift to glycolysis and increased glutamine metabolism (51, 89) to boost Treg proliferation and action. We found high levels of TNFR2 on over 60% of SF Treg clusters and the ligand TNF-α is increased in the inflamed joint (4), thus it remains to be seen which metabolic pathway Tregs utilise in SF and how this alters their functionality.

In conventional T cells, TNFR2 expression and subsequent glutamine catabolism and glycolysis have been linked to pro-inflammatory actions and impact on the Th17-Treg balance (51, 90, 91), a key imbalance implicated in the pathogenesis of JIA (19, 84). Moreover, TNFR2 expression on CD4+ conventional T cells might contribute to resistance to Treg suppression (49, 50). Similarly, resistance of JIA effector T cells to suppression by Tregs was linked to TNF-α (14, 15), although TNFR2 was not assessed in these studies. Here, a significant TNFR2high cluster (CD3+ CD4+TNFR2highPD-1low) was increased in SF conventional T cells, suggesting TNFR2, glutamine catabolism and glycolysis could be a potential target in the JIA joint to restore the immunoregulatory balance.

Despite increased presence of CD4+Foxp3+ Tregs, it has been previously suggested that the inflammatory environment in the joint can alter Treg phenotype and possible function (5). High-dimensional clustering exposed new functionally distinct Treg subsets in the inflamed site of JIA. The five SF-predominant Treg clusters were highly activated, with the highest expression of classical functional Treg markers such as CTLA-4, TNFR2 and TIGIT. The co-inhibitory ectonucleotidase CD39, however, had more varied expression across these subsets. Despite previous evidence of a large proportion of SF Tregs being enriched for CD39 (92), we identified over 40% of SF Tregs (clusters 4,9-10) to be CD39low. Since high CD39 expression has been associated with Treg stability and suppressive ability even under inflammatory conditions (54), these data question the efficient functionality of these SF Tregs. Loss of CD39 from Tregs can downregulate Foxp3 expression and indeed we demonstrated possible unstable Foxp3 expression and high PD-1 expression in Treg cluster 10, which could represent loss of IL-2 sensitivity (13, 54). Interestingly, Treg clusters 9 and 10 (both CD39low) were also the only SF-dominant clusters found enriched in blood of individuals with active joint inflammation.

SF Treg suppression assays have resulted in conflicting data on their functionality in JIA (5). The heterogeneity demonstrated here and in other recent studies (93) suggests that specific Treg subsets, with their unique receptor combination, likely have different functions. Thus, it will be important to design assays that test specific molecules’ roles in Treg fitness in accordance with the ligands/substrates found in the microenvironment of the inflamed joint.

Here, we also identified a significant cluster of CD4- T cells exclusive to the inflamed joint that expressed the transcription factor Foxp3. CD8+Foxp3+ T cells have been identified in cancer and other inflammatory settings, including in the joints of ankylosing and psoriatic arthritis (43, 44, 94, 95), but have yet to be fully defined in JIA. Similarly, this JIA SF cluster also highly expressed other Treg markers, including Helios and TIGIT. Treg-like CD8+ cells have demonstrated suppressive activity with additional cytotoxic profile compared to their CD4+ counterparts (44, 96) and in mouse models enhance therapeutic potential in combination with CD4+ Tregs (96–99). Alternatively, Foxp3 expression in CD8+ T cells has more recently been linked with metabolic reprogramming in the tumour microenvironment to promote sustained survival under restricted glucose availability, but with no suppressive capabilities (45). Whether these identified CD4-Foxp3+ T cells in the inflamed JIA joint are synergistic suppressors or a demonstration of effector T cell metabolic adaptation still needs to be determined.

CD69 was a marker unifying lymphocytes in SF. Besides being an early activation marker (33), CD69 has been linked to tissue residency (35). While SF in health is devoid of immune cells, immune infiltration occurs in active JIA and thus CD69 expression could mark immune cells for retention once within the joint, where they perpetuate local inflammation. Moreover, we show that despite hyperactivity most SF T cells were not proliferative, aligning with previous reports of a lack of T cells within S phase in JIA SF (100).

By comparing PB samples from individuals with active joint inflammation at time of sample collection to those without, we aimed to assess whether SF-predominant cell clusters were detectable as they recirculate through the blood. We found limited recirculation of subpopulations of APCs, NK and CD3 effector cells. Only SF T cell cluster 4, an exhausted-like CD4- subset, could be found at statistically significantly increased but minute frequency in blood of active compared to inactive JIA samples. Similarly, Spreafico *et. Al*. found a small population of CD4+ T cells in JIA blood that shared a similar signature and TCR clonality with paired SF T cells (101). These data, together with our data demonstrating minimal proliferation of T cells in SF, and data on homing chemokine receptors (4, 101), further support enhanced infiltration into the joint, adaptation at the site and retention.

More differences were seen between active and inactive JIA PB Tregs. Of particular interest were clusters 9 and 10 (CD71highCD39low), which made up over 30% of SF Tregs. The PB counterparts mirrored the SF Treg phenotype, thus suggesting these populations might be recirculating. These identified Tregs could be circulating to other joints and associated lymph nodes and fail to stop APCs promoting autoreactive T cell activation, which may explain worsening of disease with each flare (6).

Spectral flow cytometry allowed the assessment of phenotypes in the inflamed joint of JIA by a wider array of markers than collective conventional approaches used before. However, this study was still limited by pre-selecting which proteins to screen for, and additional cellular adaptation are likely to occur in the inflamed joint which we have yet to characterise. Fully unbiased and comprehensive approaches, such as single cell RNA sequencing, will likely reveal further complex interactions and novel cell populations. Although RNA abundance does not necessarily mirror protein abondance or functionality. Furthermore, here we exclusively investigated phenotypic states, and further studies, which were beyond the scope of this study, are required to understand the functional role of each subset. Here, we offer a methodology for identifying multiple cellular states and clusters of interest on a protein-level which can be more easily classified for further functional investigation and possible disease monitoring and therapeutic targeting. Using a cross sectional patient cohort, we investigated an overall snapshot in time of the inflamed site and peripheral blood of JIA.

## Conclusion

This is the first study assessing immune landscape by high-dimensional protein expression in each sample of cross-sectional JIA cohorts, including synovial fluid and blood of individuals with active JIA, and blood of individuals with inactive JIA and healthy controls. We characterise the immune landscape and heterogeneity in synovial fluid at a detailed high-dimensional protein level and reveal new insights into the signatures that drive joint infiltration, adaptations and possible retention of immune cell subtypes. These findings will give rise to further functional analysis of the altered immune phenotypes found exclusively in the inflamed joint. This study represents a valuable dataset and provides a platform for further functional studies, as well as cross-cohort and cross-disease comparisons. This could lead to the identification of more targeted therapies that act locally within the inflamed joint with less systemic side effects. Recirculation of minor populations of dysfunctional Tregs in active disease could potentially cause loss of immune regulation at distant sites permitting immune cell recruitment to new joints during flares. Ultimately, therapeutic targeting of the SF-specific inflammatory cells could stop perpetual inflammation and together with boosting immunoregulatory cells systemically, remission might be achieved in more individuals with JIA.

## Methods

### Sample processing and demographics

With informed consent/assent peripheral blood (PB) was collected by venepuncture and unpaired synovial fluid (SF) was collected at time of therapeutic joint aspiration prior to therapeutic intra articular joint injection. Hyaluronidase at 1 μL/mL was added to SF samples (30 mins at 37°C), before SFMCs and PBMCs isolation via density gradient centrifugation according to standard protocols and cryopreserved.

Routine clinical data including disease duration, JIA subtype, ANA, RF status and medication were extracted from the study databases or clinical records in fully anonymised fashion. Clinical Juvenile Arthritis Disease Activity Score (cJADAS) was calculated (102). Clinically inactive disease was defined as having no active joints and active disease having an active joint count (AJC) of 1 or above. Table 1 displays the demographics and clinical characteristics at the time of sample collection. For demographics and clinical characteristics for active and inactive samples see Table S2 in supplemental material.

### Flow cytometry

To comprehensively assess differences in cellular composition between PB and SF, cryopreserved samples were thawed and stained with fixable viability dye (FVD) and antibodies for surface markers CD3, CD4, CD11c, CD14, CD16, CD19, CD25, CD39, CD45RA, CD56, CD69, CD71, CD96, CD112, TNFR2, CD123, CD127, CD155, CD161, CD226, GARP, GITR, HLA-DR, Lap, PD-1, TIGIT, 4-1BB. Cells were fixed and permeabilized using Foxp3 Transcription Factor Fixation/Permeabilization buffer (eBioscience™) per the manufacturer’s instructions and stained with antibodies for intracellular markers (CTLA-4, Foxp3, Helios, ID2, Ki67). For details of the antibodies see Table S1 in supplemental material. Samples were acquired on a 5-laser Aurora (Cytek®), a full-spectrum flow cytometer. Autofluorescence (AF) was acquired as a separate parameter for AF extraction (68). Optimisation included generation of a skeleton panel, assessing fluorescence minus one (FMO) controls on *ex vivo* and *in vitro* stimulated PBMCs. SFMCs were also tested with FMOs due to known changes in cell size, autofluorescence and expression levels of certain markers, and to ensure accurate assessment of markers across the expression spectrum expected and sample sources utilised in this study. Spectral flow cytometry data is available at www.flowrepository.org under FLOWRepository ID FR-FCM-Z6VC.

### Analysis

Data was analysed using FlowJo v10 (BD Bioscience) for manual gating. For unbiased high dimensional analysis, data was analysed using Spectre (23) on R, using clustering algorithms FlowSOM or PhenoGraph, dimensionality reduction UMAP gated on single, live (FVD-) cells, debris removed, with/without specific lineage definition. Statistical analyses were performed using Graphpad Prism v10.0.2 using non-parametric t test (Mann-Whitney) or one-way ANOVA with Tukey’s multiple comparison testing and p values <0.05 were considered statistically significant.

### Study approval

This research was conducted under the informed consent of participants, in accordance with the following research ethics committees: NHS London - Bloomsbury Research Ethics Committee REC references JIAP-95RU04 and CHARMS-05/Q0508/95 studies (Wedderburn); UCL research Ethics 14017/001 and 14017/002 (Pesenacker). JIA patients were recruited to studies with full informed parental consent and age-appropriate assent (or consent for those over 16 years of age). HC PB donors were recruited with full consent for use of their blood in research.

## Author contributions

MHA contributed to the overall study design, execution of all experiments, data analysis, and writing and editing of the manuscript. DS performed data analysis, figure generation and contributed to writing and editing of the manuscript. VA, MK assisted with sample collection, processing, clinical data queries and editing of the manuscript. JIA samples were part of the CHARMS and JIAP studies. LRW oversaw subject recruitment, sample biobanking and patient data collection and contributed to the study design and editing of the manuscript. AMP led and oversaw the overall study design, acquisition, analysis and writing and editing of the manuscript, and takes responsibility for the content of the article. All authors read and approved the final version of the manuscript.

## Acknowledgments

This project was funded by NIHR BRC UCLH BRC764/III/AP/101350, and ARUK (now Versus Arthritis) CDF 21738, Versus Arthritis 23159, UKRI BBSRC BB/V009524/1 supported salaries of AMP, MHA, DS and contributed to general shared lab consumables. The work was also supported by Versus Arthritis 20164, 21593 and 22084, GOSH Children’s Charity (VS0518) and UKRI MRC CLUSTER-JIA award (MR/R013926/1), which supported salaries of MK and VA. The CLUSTER-JIA MRC award provided support for recruitment and sample cohorts; CLUSTER is additionally supported by funding, or funding in kind, from Pfizer, UCB, AbbVie, SOBI and GSK, but those parties played no role in this work. LRW and the Wedderburn lab are supported by the NIHR Biomedical Research Centre (BRC) at Great Ormond Street Hospital.

The authors would like to thank all the patients, parents, clinical staff, and study coordinators at Great Ormond Street Hospital for their contribution to this research. We acknowledge and thank Janani Sivakumaran Nguyen at the IIT flow cytometry facility for all her technical help and support. We would like to thank Dr Lizzy Rosser, Dr Laura Pallet, Prof David Sansom and Prof Lucy Walker for helpful discussions and reading of the manuscript. The authors would like to thank the teams at UCL IIT and UCL GOS ICH for all the discussion, feedback, and exchange of ideas.

## Notes

Conflict of interest statement: MHA, DS and AMP have no conflict of interest to declare. LW, MK and VA received funding, or funding in kind to the CLUSTER-JIA Consortium from Pfizer, UCB, AbbVie, SOBI and GSK, but that support did not directly fund this work. LW has received speaker fees, paid to UCL, from Pfizer, unrelated to this work.

http://flowrepository.org

### Competing Interest Statement

MHA, DS and AMP have no conflict of interest to declare.
LW, MK and VA received funding, or funding in kind to the CLUSTER-JIA Consortium from Pfizer, UCB, AbbVie, SOBI and GSK, but that support did not directly fund this work. LW has received speaker fees, paid to UCL, from Pfizer, unrelated to this work.

