## supplemental material for "The immune landscape of the inflamed joint defined by spectral flow cytometry"

**Table S1. Antibodies used in high dimensional flow cytometry panel for cellular composition and activation, proliferative and functional state of mononuclear cells**; *Antibodies used at appropriately tested titrations.*

| Fluorophore | Marker | Clone |
|---|---|---|
| BUV395 | CD4 | SK3 |
| BUV496 | HLA-DR | G46-6 |
| BUV563 | CD161 | DX12 |
| BUV615 | CD16 | 3G8 |
| BUV661 | 4-1BB (CD137) | 4B4-1 |
| BUV737 | CD25 | 2A3 |
| BUV805 | CD39 | TU66 |
| BV421 | TNFR2 (CD120b) | hTNFR-M1 |
| SB436 | CD123 | 6H6 |
| eFluor450 | ID2* | ILCID2 |
| BV480 | CD127 | HIL-7R-M21 |
| BV510 | CD71 | M-A712 |
| BV570 | CD56 | 5.1H11 |
| BV605 | CTLA-4*(CD152) | BNI3 |
| BV650 | Ki67* | B56 |
| BV711 | CD226 (DNAM-1) | 11A8 |
| BV750 | GARP | 7B11 |
| BV785 | CD69 | FN50 |
| Qdot 800 | CD14 | TuK4 |
| BB515 | CD96 | 6F9 |
| AF488 | GITR (CD357) | 108-17 |
| AF532 | CD45RA | HI100 |
| BB700 | PD-1 (CD279) | EH12.1 |
| PerCP-eFluor710 | Lap | FNLAP |
| PE | Foxp3* | 236A/E7 |
| PE-CF594 | CD11c | 3.9 |
| PE-Cy5 | CD19 | HIB19 |
| PC-Cy7 | CD112 | TX31 |
| APC | CD155 (PVR) | SKII.4 |
| AF647 | TIGIT | MBSA43 |
| AF700 | Helios* | 22F6 |
| eFluor 780 | FVD | --- |
| APC-Fire810 | CD3 | SK7 |

**stained intracellularly after FoxP3/transcription factor fixation/permeabilization buffer*

**Table S2. JIA PB cohort divided by clinically active or inactive disease**

Of all JIA PBMC samples, 'Active' Juvenile Idiopathic Arthritis (JIA) PB classified by one or more clinically active joint; 'Inactive' JIA PB classified by no clinically active joints at time of sample of those where AJC data was available. Shown as mean (range) unless otherwise specified.

| | JIA PBMC 'Inactive' | JIA PBMC 'Active' |
|---|---|---|
| **Number of participants, n** | 17 | 29 |
| **Gender, % Female** | 88.2 | 65.5 |
| **Age at sample in years, mean (range)** | 9.6 (2-18) | 5.7 (1-16) |
| **Ethnicity, % Caucasian** | 82.3 | 62.1 |
| **Disease duration:** | | |
| **Time since diagnosis, months (range)** ◊ | 47.1 (3-125) | 31.6 (2-105) |
| **JIA subtype:** | | |
| **% RF- Polyarticular** | 64.7 | 21.4 |
| **% Oligoarticular** | 35.3 | 78.6 |
| **Medication at time of sample:** | | |
| **Methotrexate, n** | Yes= 11<br>No= 6 | Yes= 15<br>No= 14 |
| **Steroids, n** ∇ | Yes= 10<br>No= 7 | Yes= 13<br>No= 15 |
| **Biologics, n** | Yes= 1<br>No= 16 | Yes= 3<br>No= 26 |
| **Clinical information at time of sample:** | | |
| **ANA, n** # | positive = 7<br>negative = 4 | positive = 21<br>negative = 6 |
| **AJC, mean (range)** | 0 (0) | 2.03 (1-5) |
| **cJADAS, mean (range)** ⅄ | 1.8 (0-8.6) | 7.9 (1.7-17.3) |

PBMC= Peripheral Blood Mononuclear Cells; RF-: Rheumatoid factor negative; ANA: Antinuclear Antibodies (positive classified as titre ≥1:160); AJC: Active Joint Count; cJADAS: clinical Juvenile Arthritis Disease Activity Score

◊ 5.9% of Inactive JIA PBMC samples missing data; ∇ 2.4% of Active JIA PBMC samples missing data; **#** 35.2% of Inactive JIA PBMC and 6.9% of Active JIA PBMC samples missing data; ♦11.5% of JIA PBMC samples missing data; ⅄11.8% of Inactive JIA PBMC and 13.8% of Active JIA PBMC samples missing data
